# Ultrasound-based optimal parameter estimation improves assessment of calf muscle-tendon interaction during walking

**DOI:** 10.1101/778720

**Authors:** T. Delabastita, M. Afschrift, B. Vanwanseele, F. De Groote

## Abstract

We present and evaluate a new approach to estimate calf muscle-tendon parameters and calculate calf muscle-tendon function during walking. We used motion analysis, ultrasound, and EMG data of the calf muscles collected in six young and six older adults during treadmill walking as inputs to a new optimal estimation algorithm. We used estimated parameters or scaled generic parameters in an existing approach to calculate muscle fiber lengths and activations. We calculated the fit with experimental data in terms of root mean squared differences (RMSD) and coefficients of determination (R^2^). We also calculated the calf muscle metabolic energy cost. RMSD between measured and calculated fiber lengths and activations decreased and R^2^ increased when estimating parameters compared to using scaled generic parameters. Moreover, R^2^ between measured and calculated gastrocnemius medialis fiber length and soleus activations increased by 19 % and 70 %, and calf muscle metabolic energy decreased by 25% when using estimated parameters compared to using scaled generic parameters at speeds not used for estimation. This new approach estimates calf muscle-tendon parameters in good accordance with values reported in literature. The approach improves calculations of calf muscle-tendon interaction during walking and highlights the importance of individualizing calf muscle-tendon parameters.

## Introduction

The calf muscles are important for forward propulsion during human walking ^26^. These muscles can efficiently produce high forces during push-off thanks to the muscle-tendon interactions. The Achilles tendon elastic properties decouple calf muscle fiber length from calf muscle-tendon unit length during walking ^20^. In young adults, Achilles tendon stiffness seems to be optimized to enable the calf muscle fibers to work nearly isometric and near their optimal fiber length ^20,22^, i.e., the condition that allows maximal force output. In older adults, Achilles tendon stiffness is decreased ^6^ and the Achilles tendon lengthens more during the stance phase of walking ^23^. Hence, the calf muscle fibers work at higher contraction velocities and possibly lower length ^23^. It has been suggested that these less optimal muscle working conditions increase the activated muscle volume and hence the metabolic energy needed to produce the same forces ^22^. However, studying the contribution of decreased Achilles tendon stiffness to increased metabolic cost of walking experimentally is hard due to the many simultaneous age-related changes in the musculoskeletal system. Musculoskeletal modelling and simulations might reveal such causal relationships by allowing us to compute the isolated effect of changes in Achilles tendon stiffness on calf muscle metabolic cost based on simulated fiber lengths, activations and forces.

Investigating the influence of elastic tendon properties on muscle activations, fiber lengths, and forces requires simulation methods that account for muscle-tendon interactions. Commonly, optimization (minimizing a performance criterion e.g. muscle activations squared) is used to compute muscle excitations from measured joint kinematics and joint moments. Specifically, static optimization has mostly been applied ^25^ but this approach assumes rigid tendons ^5^. However, we recently developed a numerically efficient dynamic optimization approach that accounts for the influence of the elastic tendon on the muscle fiber’s working length and velocity ^15^. This new approach is suitable to investigate the influence of Achilles tendon stiffness on muscle-tendon function during walking.

Investigating calf muscle-tendon interaction and its effect on metabolic cost of walking for individual subjects requires personalization of the calf-muscle tendon properties in the model. Muscle-tendon properties vary greatly with age, gender and physical activity level ^33^ and simulation outputs are very sensitive to variations in optimal muscle fiber length, tendon slack length, and tendon stiffness ^14,22^.

B-mode ultrasound provides information about muscle-tendon interaction that could be used to individualize calf muscle-tendon properties in musculoskeletal models ^27^ but existing approaches have not been validated. In previous research, ultrasound-based gastrocnemius medialis muscle architecture and the Achilles tendon force-strain relationship during an isometric contraction were used to individualize musculoskeletal models and these models were used to simulate gastrocnemius medialis fiber length during isometric contractions, running and hopping ^12,13^. However, the authors did not validate the simulated gastrocnemius medialis fiber length with experimental data in these dynamic tasks.

Our aim was therefore to develop and validate a new dynamic optimization approach to individualize calf muscle-tendon parameters based on information from B-mode ultrasound images collected during walking. We hypothesized that the use of individualized parameters would improve the accuracy of calculated calf muscle activations and fiber lengths at walking speeds different from the speeds used to individualize the parameters. In addition, we hypothesized that the use of personalized instead of scaled generic calf muscle-tendon properties would greatly influence the computation of metabolic energy.

## Materials and Methods

### Experimental data collection

Six healthy young adults (mean age 24.5 ± 2.7 years, height 169.5 ± 8.6 cm; body mass 62.4 ± 7.5 kg; two male and four female) and six healthy older adults (mean age 77.1 ± 5.0 years, height 165.7 ± 9.2 cm; body mass 71.4 ± 9.9 kg; three male and three female) were included in this study. We excluded subjects that participated in competitive sports or had exercised more than twice a week in the previous three months. The Medical Ethics Committee UZ/KU Leuven approved the study and subjects signed written consent prior to participation in accordance with the Declaration of Helsinki.

First, we determined the subject’s comfortable and maximal overground walking speeds. We calculated their preferred speed as the time to cover ten meters that were part of a demarcated track. The subjects were asked to walk at a comfortable speed that can easily be maintained. We calculated their maximal walking speed as the mean speed during a 6-minute walking test. Next, participants familiarized with the instrumented treadmill (Motekforce Link, Amsterdam, The Netherlands) for minimally 20 minutes. During treadmill testing, subjects walked at their comfortable and maximal walking speeds, at 3 km/h and, at 5 km/h. Between the walking speeds, patients were allowed to sit until they indicated that they were sufficiently recovered. For further analysis, we ordered the speeds in increasing order. We refer to the speeds by speed 1, 2, 3, and 4, because the order was subject-specific. We collected motion capture data, ultrasound images, and EMG signals simultaneously for five consecutive strides after 5 to 8 minutes of adaptation to the treadmill at each walking speed.

We captured the trajectories of 68 skin mounted reflective markers (extended full-body plug-in-gait marker set including nine cluster markers ^30^) using a thirteen-camera motion capture system (Vicon, Oxford, UK) at 100 Hz.

Images of the gastrocnemius medialis muscle fibers of the left leg were collected at 60 Hz using a PC-based ultrasound system (Echoblaster 128, UAB Telemed, Vilnius, Lithuania). The flat ultrasound probe with a wave frequency of 8 MHz and a field of view of 60 mm was carefully placed in line with the muscle fibers and perpendicular to the deep aponeurosis in the middle of the gastrocnemius medialis muscle belly ^3^ and attached using a custom-made plastic device and bandage^21^.

Electromyography (EMG) signals of the gastrocnemius lateralis, soleus and tibialis anterior of the right leg were collected at 1000 Hz (ZeroWire EMG, Cometa, Milano, Italy).

### Data processing

OpenSim’s gait2392-model (5 degrees of freedom and 43 muscles per leg) was scaled to the subject’s anthropometry using OpenSim’s 3.2 scale tool ^7^ based on marker data collected during a static trial. Joint kinematics were calculated with a Kalman smoothing algorithm for inverse kinematics ^16^. Afterwards, joint moments, muscle tendon lengths and muscle moment arms were calculated with OpenSim’s inverse dynamics and muscle analysis tools ^7^.

Ultrasound images were processed by tracking three lines representing the superficial aponeurosis, the deep aponeurosis and the orientation of the muscle fibers using a semi-automated algorithm ^8^. We linearly extrapolated these lines. We defined muscle fascicle length as the distance between the superficial and the deep aponeurosis parallel to the muscle fibers, and pennation angle as the angle between the muscle fibers and the deep aponeurosis^20^.

EMG signals were band-pass filtered (20 – 400 Hz), rectified, and low-pass filtered (10 Hz). We assumed that EMG signals from the muscles of the right leg were a good approximation for the EMG signals from the muscles of the left leg. We calculated the mean EMG signal from the muscles of the right leg over the recorded strides from right heel strike to right heel strike. These mean EMG signals were then time shifted to represent the EMG signals of the left leg over the strides included in the algorithms from left heel strike to left heel strike. To enable the use of these mean EMG signals to constrain muscle activations during the simulations, we processed this signal with a first order differential equation representing activation dynamics^15^. We further refer to these processed EMG signals by measured activations.

### Optimal parameter estimation

A previously developed dynamic optimization approach ^15^ to solve the muscle redundancy problem, i.e., to calculate muscle activations and forces underlying a measured movement while accounting for muscle dynamics, was modified to estimate calf muscle-tendon parameters. We solved for muscle-tendon parameters that minimized the sum of squared muscle activations as well as the difference between simulated and measured gastrocnemius medialis fiber length and pennation angle, and muscle activations patterns from the gastrocnemius lateralis, soleus, and tibialis anterior muscles.

#### Static parameters

We estimated Achilles tendon stiffness, gastrocnemius medialis and lateralis optimal fiber lengths, tendon slack lengths, and gastrocnemius medialis pennation angle at optimal fiber length. Previous research showed either large inter-subject variability of these parameters ^33^ or a large influence of these parameters on the solution of the muscle redundancy problem ^14,22^. Gastrocnemius medialis pennation angle was included because we observed important differences between simulated and measured pennation angles when piloting the study. In addition, we added three static parameters to impose the measured muscle activations in the simulations. These parameters represent the scaling of the EMG signal to simulated muscle activity of the gastrocnemius lateralis, soleus, and tibialis anterior. As such, we imposed the pattern and not the amplitude of the measured muscle activations.

#### Cost functional

The cost functional consisted of three parts and minimizes

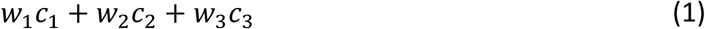

where *w*_1_ = 1, *w*_2_ = 20, and *w*_3_ = 0.5 are weight factors for the different parts of the cost functional. To select the weights on the different parts in the cost function, we visualized the influence of increasing and decreasing the weights of one term on the other tracking terms in the cost function. We chose the weight that reduced the term of interest but did not increase the other terms in the cost function (Pareto optimality ^31^).

The first part consisted of muscle and reserve actuator activations, *a*_*m*_ and *a*_*T,K*_ respectively^15,17^:

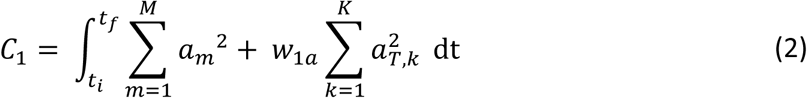

where *t*_*i*_ and *t*_*f*_ are the initial and final times; m = 1, … M indicates the muscles in the model; *w*_1*a*_ = 1000 is a weight factor; k =1, … K indicates the degrees of freedom in the model. The second part penalized differences between measured and simulated gastrocnemius medialis fiber lengths, 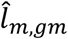 and *l*_*m,gm*_ respectively, and pennation angles, 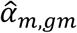 and *α*_*m,gm*_ respectively:

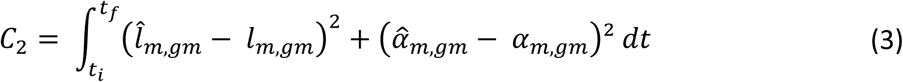

where *w*_2_= 20 is a weight factor. For one subject *w*_2_= 10 because using *w*_2_= 20 led to convergence issues. The third part penalized differences between measured and calculated muscle activations, â_*m*_ and *a*_*m*_ respectively:

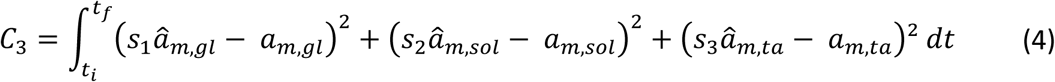

where *w*_3_= 0.5 is a weight factor; *s*_1_, *s*_2_, and *s*_3_ are the static parameters used to scale the measured muscle activations; *gl, sol*, and *ta* refer to gastrocnemius lateralis, soleus, and tibialis anterior respectively. Including the measured muscle activations in the cost functional allowed slight deviations from the measured activations to account for possible differences between left and right muscle activations at higher walking speeds ^24^. For two subjects, measured muscle activations for the gastrocnemius lateralis were not tracked because the EMG signals were not available at all walking speeds.

#### Constraints

The sum of the three parts *C*_1_, *C*_2_, and *C*_3_ was minimized subject to muscle activation and contraction dynamics ^15^. In contrast to the muscle-tendon dynamics described previously ^15,17^, tendons were modelled as linear springs ^22^.

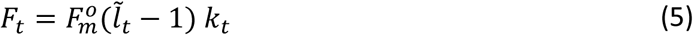

where *F*_*t*_ is tendon force,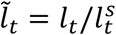 is normalized tendon length, and *k*_*t*_ is normalized tendon stiffness. The sum of the muscle torques was constrained equal to the inverse dynamics torques for every degree of freedom ^15^.

We bounded the static parameters based on ultrasound data or within physiologically realistic limits. Previous research suggests that gastrocnemius medialis muscle fibers act close to their optimal fiber length during the nearly isometric stance phase of walking ^11^. Therefore, the upper and lower bound of gastrocnemius medialis optimal fiber length were set to the maximal measured value and to this value decreased with 40% of the measured range, respectively. We imposed the following bounds:

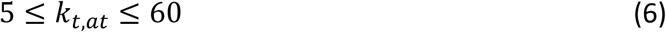

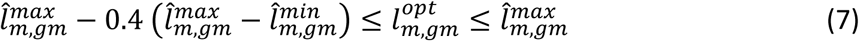

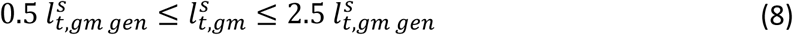

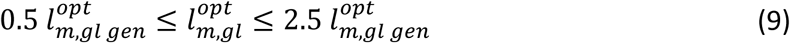

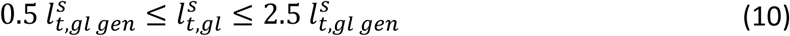

where *k*_*t,at*_ is Achilles tendon stiffness; 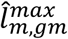, and 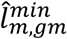 are the maximal and minimal values for gastrocnemius medialis fiber length measured during the strides included in the estimation problem; 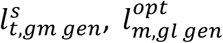, and 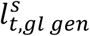 are the values in the scaled generic model for respectively gastrocnemius medialis tendon slack length 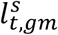, and gastrocnemius lateralis optimal fiber length 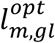 and tendon slack length 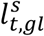. Additional constraints were added to enforce that the gastrocnemius medialis and lateralis operate in the same range of the force-length curve ^4^ and have comparable activation patterns ^35^.

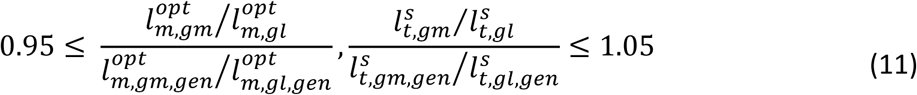

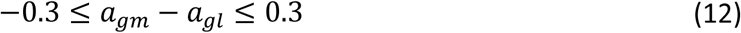

where *a*_*gm*_ and *a*_*gl*_ are the calculated muscle activations for gastrocnemius medialis and lateralis, respectively.

#### Solving the optimal parameter estimations

We applied this new algorithm to two strides of walking at speed 1, two strides of walking at speed 3, or two strides of walking at speed 1 combined with two strides of walking at speed 3 resulting in three sets of estimated calf muscle-tendon parameters. Ideally, the algorithm estimates the same parameters independent of the speed of the strides included in the algorithm. Each stride represented a different phase in one optimal parameter estimation problem. These multiple-phase problems were solved with direct collocation using GPOPS ^29^, ADiGator ^28^ and Ipopt ^34^ as described previously ^15^.

### Evaluation of the estimated parameters

We evaluated our approach in three steps. First, we evaluated the fit between measured and outputs after parameter optimization at speeds 1 and/or 3 (Fig. 1, upper left side, yellow rectangles) to assess to what extent the proposed model and optimized parameters can describe the measured outputs. We refer to these outputs after parameter optimization as estimated outputs. To this aim, we calculated the root mean squared difference (RMSD) and the coefficient of determination (R^2^) between estimated and measured gastrocnemius medialis fiber length and pennation angle, as well as R^2^ between estimated and measured gastrocnemius lateralis, soleus, and tibialis anterior muscle activations. We did not compute the RMSD between simulated and measured muscle activations since EMG does not provide information about signal amplitude. We compared these RMSD and R^2^ values to the corresponding RMSD and R^2^ between generic and measured outputs with dependent t-tests. We refer to these outputs from the algorithm using a model with scaled generic muscle-tendon parameters as generic outputs. The generic outputs were obtained by solving the muscle redundancy problem ^15^ based on a scaled, generic musculoskeletal model (Fig. 1, lower left side, red dashed rectangles).

**Fig. 1.**
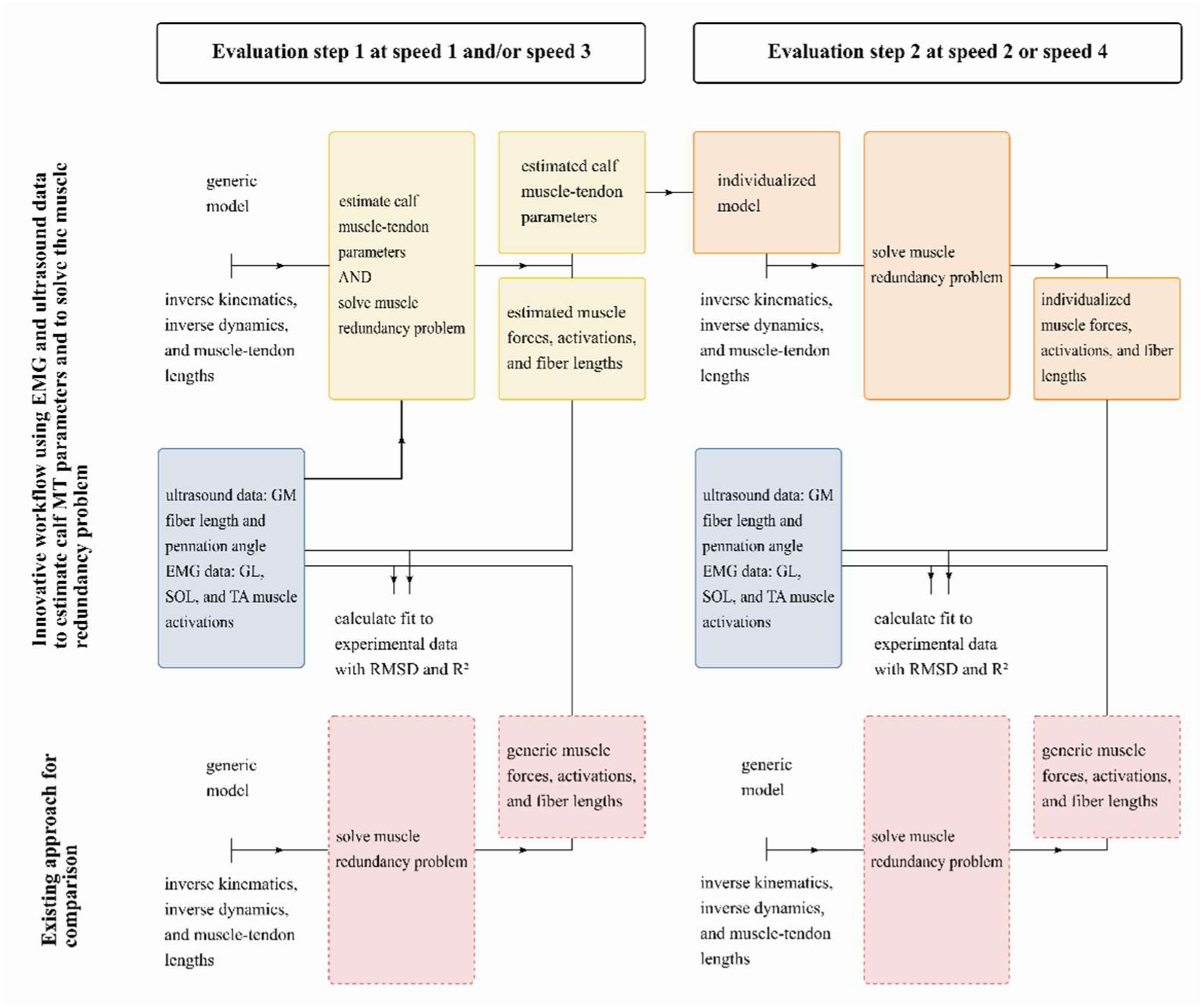
Workflow of the approach to evaluate a new method to estimate individualized calf muscle-tendon parameters and calculate muscle forces, activations and fiber lengths. Yellow rectangles present the new optimal estimation algorithm; orange rectangles present the existing algorithm that used an individualized musculoskeletal model with estimated calf muscle-tendon parameters; the red rectangles with dashed outline present the existing algorithm that used a (scaled) generic musculoskeletal model; blue rectangles present the measured experimental data used as input in the new optimal estimation algorithm or for comparison elsewhere; we performed these comparisons by calculating the RMSD and R^2^ between measured and estimated (yellow), individualized (orange), and generic outcomes (red) at speed 1 or/and speed 3 (left part), and at speed 2 and 4 (right part). All subjects’ data and models were included in all these evaluation steps. GM = gastrocnemius medialis; GL = gastrocnemius lateralis; SOL = soleus; TA = tibialis anterior; EMG = electromyography.

Second, we evaluated the fit between measured and outputs after individualizing the calf muscle-tendon parameters using the parameters estimated at speeds 1 and/or 3 to compute muscle activations and fiber lengths at speeds 2 and 4 (Fig. 1, upper right side, orange rectangles). We refer to the outputs of the approach using a model with individualized calf muscle-tendon parameters as individualized outputs. We assessed if the fit between measured and individualized outputs was better than the fit between measured and generic outputs by also using scaled generic parameters to compute muscle activations and fiber lengths at speeds 2 and 4 (Fig. 1, lower right side, red dashed rectangles). Speed 2 was between the speeds used for parameter estimation, whereas speed 4 was higher than the speeds used for parameter estimation. This allowed us to assess whether we successfully identified individualized parameters as opposed to merely fitting the experimental data. Again, we calculated the RMSD and the R^2^ between the estimated and measured values. We used dependent t-tests to compare individualized and generic RMSD and R^2^ values.

Third, we evaluated if the estimated parameters were consistent across estimations based on different input data (speed 1, speed 3, or speed 1 and 3) and in agreement with previously reported calf muscle-tendon parameters from experimental studies. We assessed the consistency of the estimated parameters based on the range of estimated parameters across estimation speeds. To allow comparison with reported Achilles tendon stiffness values in literature, we de-normalized estimated normalized stiffness ^36^ by multiplying normalized stiffness by the summed soleus, gastrocnemius medialis and lateralis maximal isometric muscle forces and by dividing it by estimated gastrocnemius medialis tendon slack length.

To demonstrate the importance of using personalized muscle-tendon properties of the calf muscles for model-based estimation of the metabolic cost of walking, we evaluated the influence of individualizing these parameters on calf muscle metabolic energy consumption during walking. We computed the metabolic energy consumption rate for the main muscles around the ankle joint, i.e. gastrocnemius medialis, lateralis, soleus, and tibialis anterior, using the model proposed by Bhargava et al. ^2^. We used dependent t-tests to compare the summed metabolic energy consumption of these plantarflexors when using estimated and scaled generic parameters.

We conducted separate analyses to account for the possible confounding influence of gender on muscle-tendon parameters. Since the number of male and female participants was not equal in this study, and since muscle-tendon parameters differ between males and females, we compared the individualized RMSD and R^2^ values between male and female participants. We used independent t-tests to compare these values at speed 2 and at speed 4 seperately. Statistical differences would indicate that our new approach estimates calf muscle-tendon parameters better in males or females.

For all statistical tests, the level of statistical significance was set at *α* = 0.05. We applied bias-corrected and accelerated bootstrapping procedures to account for small sample sizes. Statistical analyses were performed in SPSS (SPSS Statistics 25.0, IBM, New York, United States).

## Results

Optimizing calf muscle-tendon properties improved the fit between estimated and experimental measures with respect to using scaled generic parameters for gastrocnemius medialis fiber length and pennation angle, and soleus and tibialis anterior activations but did not affect the fit for gastrocnemius lateralis activation (Table I and II).

**TABLE I.**
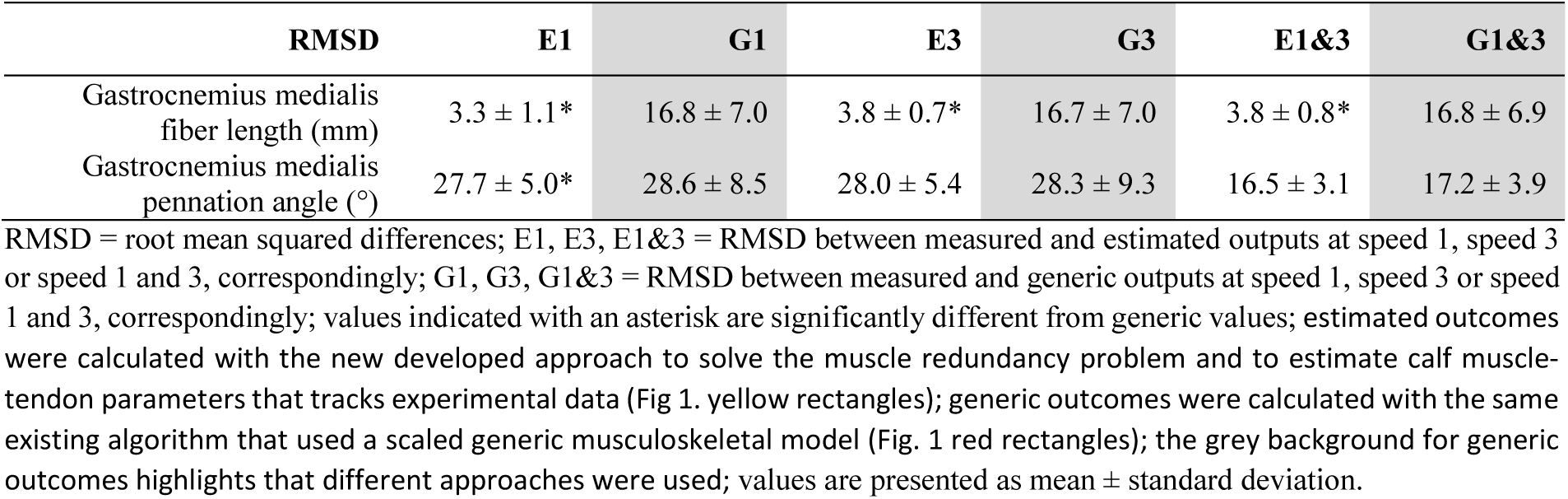
RMSD BETWEEN MEASURED AND ESTIMATED OR GENERIC OUTPUTS

**TABLE II.** R^2^ BETWEEN MEASURED AND ESTIMATED OR GENERIC OUTCOMES

| R<sup>2</sup> [ ] | E1 | G1 | E3 | G3 | E13 | G13 |
| --- | --- | --- | --- | --- | --- | --- |
| Gastrocnemius medialis fiber length | 0.71 ± 0.13* | 0.48 ± 0.20 | 0.76 ± 0.08* | 0.62 ± 0.13 | 0.73 ± 0.09* | 0.56 ± 0.14 |
| Gastrocnemius medialis pennation angle | 0.51 ± 0.25* | 0.42 ± 0.29 | 0.55 ± 0.20* | 0.47 ± 0.21 | 0.53 ± 0.19* | 0.45 ± 0.21 |
| Gastrocnemius lateralis muscle activation | 0.71 ± 0.19 | 0.60 ± 0.24 | 0.68 ± 0.29 | 0.63 ± 0.33 | 0.69 ± 0.22 | 0.61 ± 0.25 |
| Soleus muscle activation | 0.54 ± 0.23* | 0.33 ± 0.22 | 0.60 ± 0.20* | 0.28 ± 0.16 | 0.58 ± 0.17* | 0.30 ± 0.18 |
| Tibialis anterior muscle activation | 0.42 ± 0.13* | 0.31 ± 0.16 | 0.48 ± 0.22 | 0.43 ± 0.22 | 0.46 ± 0.17* | 0.37 ± 0.17 |

For gait speeds not used for parameter estimation, using the estimated parameters clearly improved the fit between simulated and experimental gastrocnemius medialis fascicle length (Table III, Fig. 2A). It did hardly alter the fit between simulated and experimental gastrocnemius medialis pennation angle (Table III, Fig. 2B) or gastrocnemius lateralis muscle activation (Fig. 2C). Moreover, it clearly improved the fit between simulated and experimental soleus muscle activation (Fig. 2D) but only slightly improved the fit between simulated and experimental tibialis anterior muscle activation (Fig. 2E). RMSD between measured and simulated gastrocnemius medialis length were 68% smaller when using individualized instead of generic parameters (Table III) and the corresponding R^2^ values were on average 19% higher (Fig. 2A). R^2^ between measured and simulated soleus activations was on average 74% higher when using individualized instead of generic parameters (Fig. 2D).

**Fig. 2.**
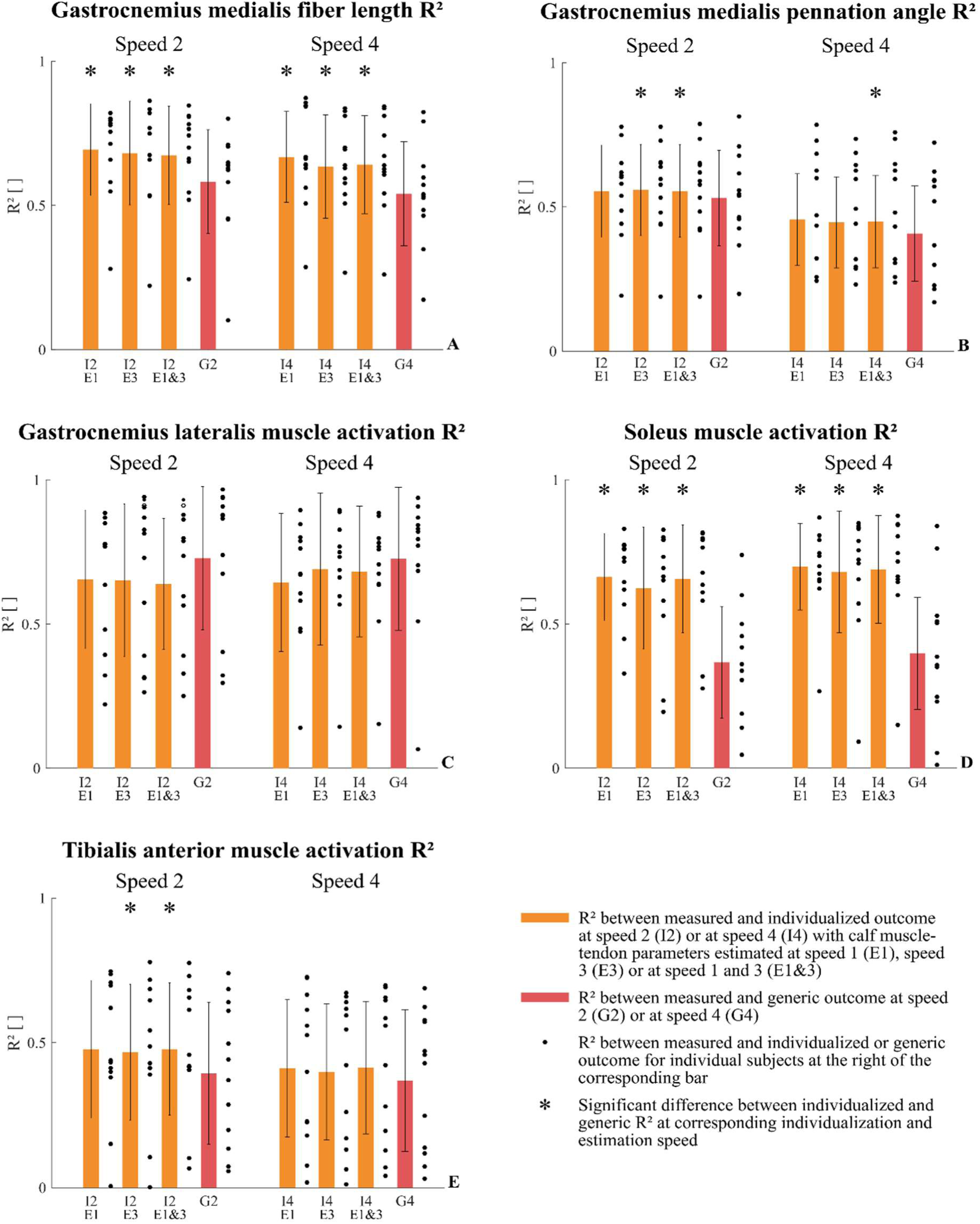
R^2^ between measured and individualized or generic outcomes. I2 or I4 = R^2^ between measured and individualized outcome at speed 2 or speed 4, correspondingly; E1, E3 or E1&3 indicate that the parameters to calculate individualized outcomes were estimated at speed 1, 3 or 1&3, correspondingly; G2 or G4 = R^2^ between measured and generic outcomes at speed 2 or speed 4, correspondingly; individualized outcomes were calculated with an existing algorithm that used an individualized musculoskeletal model with estimated calf muscle-tendon parameters (Fig.1 orange rectangles); generic outcomes were calculated with the same existing algorithm that used a scaled generic musculoskeletal model (Fig. 1 red rectangles); we explained the inter-subject variability using the results of a representative example (gastrocnemius medialis fiber length R^2^ I4 E1&3 of 0.61); we also provided individual data of a good example (gastrocnemius medialis fiber length R^2^ I4 E1&3 of 0.84) and a bad example (gastrocnemius medialis fiber length R^2^ I4 E1&3 of 0.26) in terms of the fit between individualized and measured outcomes.

**TABLE III.**
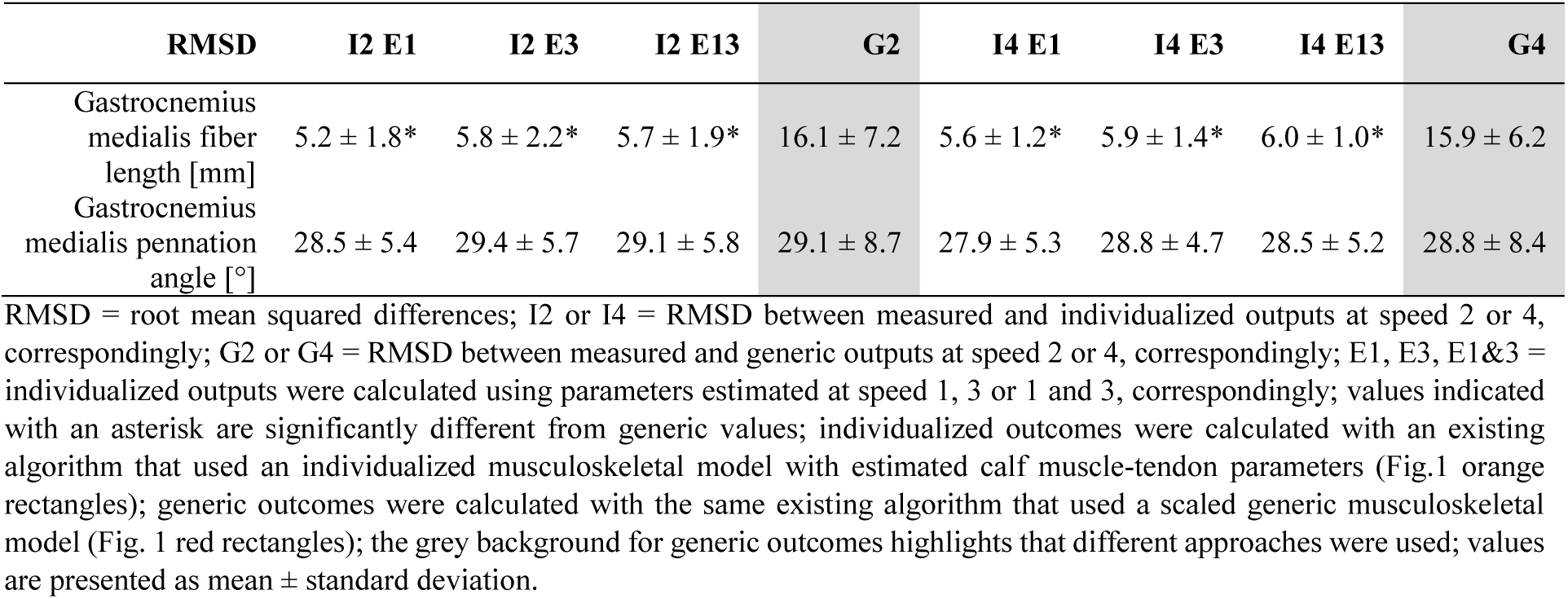
RMSD BETWEEN MEASURED AND INDIVIDUALIZED OR GENERIC OUTCOMES

The mean range across the estimates obtained with input data at different speeds was small for all parameters except normalized Achilles tendon stiffness. The mean range across estimations was 3.3±1.7 mm for gastrocnemius medialis fiber length, 3.8±1.6 mm for gastrocnemius lateralis optimal fiber, 2.8±1.4 mm for gastrocnemius medialis tendon slack length, 8.5±7.7 mm for gastrocnemius lateralis tendon slack length, and 1.0±0.8 ° for gastrocnemius medialis pennation angle at optimal fiber length. The mean range across estimations was larger for normalized Achilles tendon stiffness (3.7±2.3). Moreover, Achilles tendon stiffness estimated at speed 1 was lower than Achilles tendon stiffness estimated at speed 3 in 10 out of 12 subjects. We found no differences in estimated parameters between young and older adults (Table IV).

**TABLE IV.**
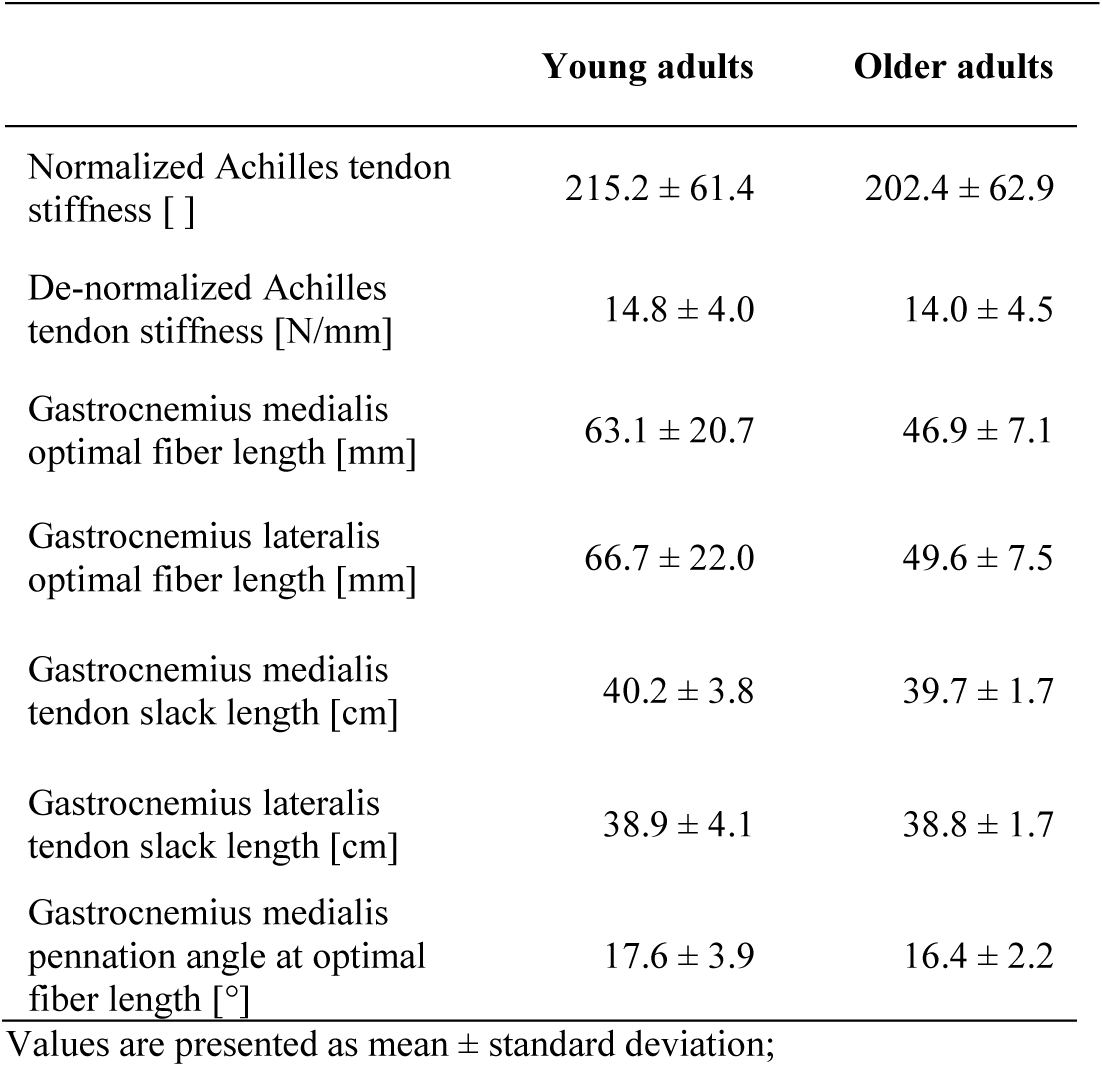
ESTIMATED PARAMETERS IN YOUNG AND OLDER ADULTS

|  | Young adults | Older adults |
| --- | --- | --- |
| Normalized Achilles tendon stiffness [ ] | $215.2 \pm 61.4$ | $202.4 \pm 62.9$ |
| De-normalized Achilles tendon stiffness [N/mm] | $14.8 \pm 4.0$ | $14.0 \pm 4.5$ |
| Gastrocnemius medialis optimal fiber length [mm] | $63.1 \pm 20.7$ | $46.9 \pm 7.1$ |
| Gastrocnemius lateralis optimal fiber length [mm] | $66.7 \pm 22.0$ | $49.6 \pm 7.5$ |
| Gastrocnemius medialis tendon slack length [cm] | $40.2 \pm 3.8$ | $39.7 \pm 1.7$ |
| Gastrocnemius lateralis tendon slack length [cm] | $38.9 \pm 4.1$ | $38.8 \pm 1.7$ |
| Gastrocnemius medialis pennation angle at optimal fiber length [°] | $17.6 \pm 3.9$ | $16.4 \pm 2.2$ |

Values are presented as mean ± standard deviation;

Using a musculoskeletal model with individualized instead of scaled generic calf muscle-tendon parameters in simulations of human walking has a big influence on calculations of calf muscle metabolic energy consumption. The average metabolic energy consumption rate across the whole stride for the triceps surae muscles was on average 25% and 35% lower when using individualized instead of generic muscle-tendon properties at speed 1 (generic 318±47 W/kg, individualized 234±33 W/kg) and at speed 3 (generic 878±227 W/kg, individualized 549±133 W/kg) respectively. This is illustrated in a representative example (Fig. 3). The influence is highest for the plantarflexors (Fig. 3A-C), and smaller for the tibialis anterior (Fig. 3D). We selected this specific representative example because the R^2^ between measured and individualized gastrocnemius medialis fiber length was 0.61, under the mean across all subjects. We added other examples (the highest R^2^ and the lowest R^2^ between measured and individualized gastrocnemius medialis fiber length, 0.84 and 0.26 correspondingly) as supplementary figures.

**Fig. 3.**
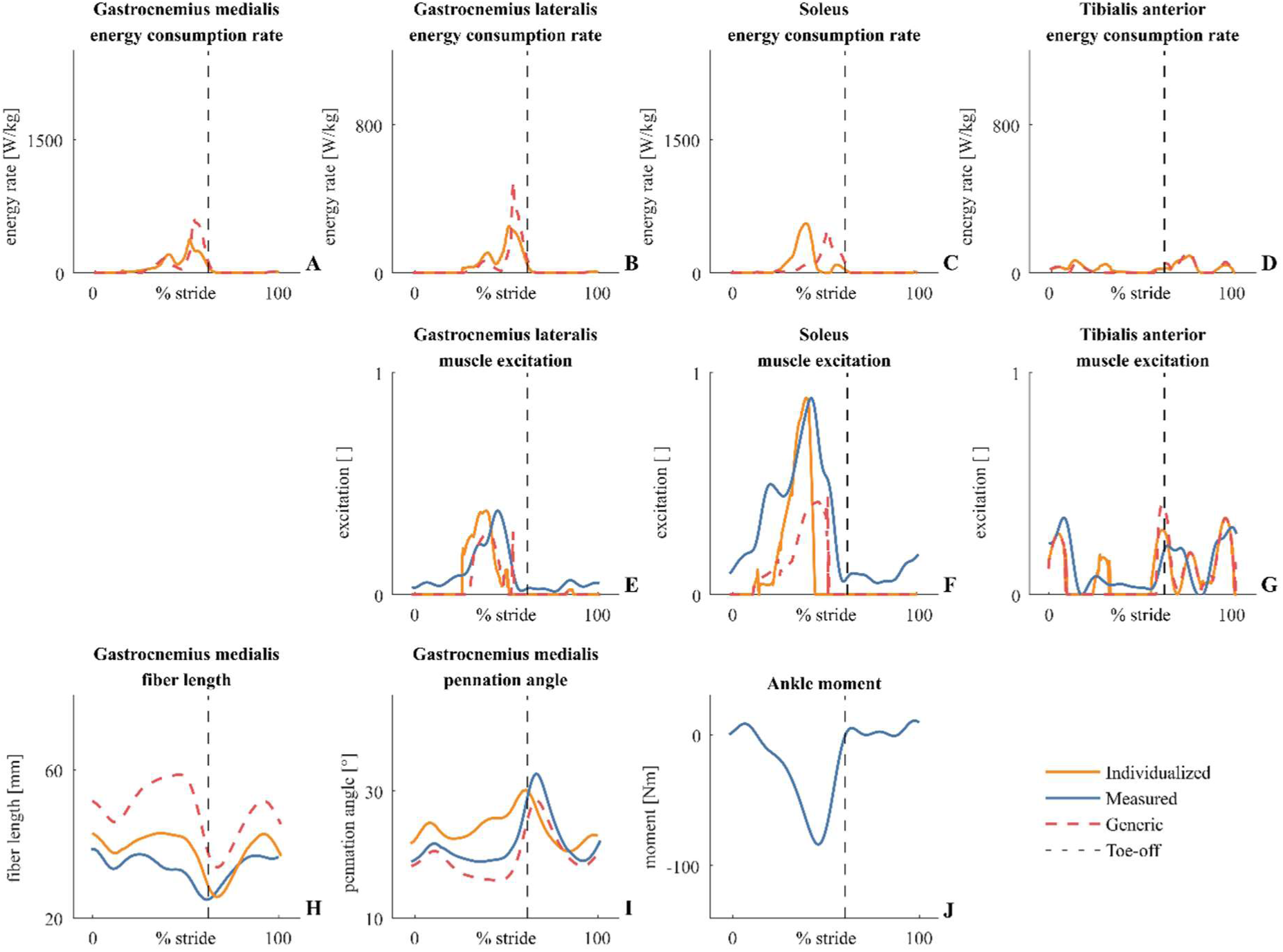
Representative example of measured, individualized, and generic outcomes for one older subject walking at 4.7 km/h with parameters estimated at speed 1 and 3. The dashed line indicates toe-off; orange lines present individualized outcomes; individualized outcomes were calculated with an existing algorithm that used an individualized musculoskeletal model with estimated calf muscle-tendon parameters (Fig.1 orange rectangles); red dashed lines present generic outcomes; generic outcomes were calculated with the same existing algorithm that used a scaled generic musculoskeletal model (Fig. 1 red rectangles); blue lines present experimental data (Fig 3. blue rectangles).

We found no statistical differences between males and females with respect to the individualized RMSD of R^2^ values. We added bar graphs for every parameter as supplementary material to illustrate the results.

## Discussion

We presented and evaluated a new approach to estimate subject-specific calf muscle-tendon parameters using ultrasound and EMG data collected during walking. Using subject-specific instead of scaled generic parameters improved calculations of calf muscle-tendon function during walking and has a large effect on calculations of metabolic energy consumption, underlining the importance of using subject-specific parameters for model-based assessment of calf muscle metabolic cost during walking. Although the use of ultrasound-derived measurements of muscle fiber lengths to estimate muscle tendon properties is not new ^12,13^, we showed for the first time that such approach improves calculations of calf muscle-tendon function during dynamic tasks such as walking.

We demonstrated that the estimated parameters because they improve the fit between simulated and measured fiber lengths and activations at walking speeds higher than the walking speeds at which we collected the data for the optimal parameter estimations. Thereby, we showed that we identified model parameters rather than fitting the data. Typically, overfitting would result in good fits between simulations and measurements when estimating subject-specific parameters but poor fits when using a model with individualized parameters to analyze data that we did not use for parameter estimation. Here, we found that using subject-specific instead of scaled generic parameters improved the computation of gastrocnemius medialis fiber length and soleus muscle activations. However, it had no or smaller influence on the fit between simulated and measured gastrocnemius lateralis and tibialis anterior activity, probably because the fit was already good for gastrocnemius lateralis and the influence of individualizing calf muscle tendon properties on tibialis anterior activity is small. We illustrated the good fit for the gastrocnemius lateralis (Fig. 3E), the improved fit for the soleus (Fig. 3F) and the poor fit for the tibialis anterior (Fig. 3G) in a representative example. These muscle activations are inter-related because together, these muscles mainly produce the muscle forces necessary to generate the ankle moments during walking. Importantly, we validated the optimal parameter estimations both at walking speeds that were in between and above the speeds used for estimation of the parameters. In contrast, Gerus et al. only validated their parameter estimations in isometric contractions or in running and hopping trials that were very similar to the trials used for estimation of the parameters ^12,13^. However, it is known from previous studies that calf muscle-tendon interaction changes with increasing walking or running speed ^19^. Furthermore, they used ultrasound data and joint moments to estimate calf muscle-tendon parameters but only joint moments to validate the parameters. Although we validated our parameter estimations at different walking speeds, care should be taken when extrapolating the results from our study to other movements. Our validation was entirely based on data collected during walking and it remains to be validated whether the subject-specific parameters also improve the estimation of muscle activations and fiber lengths for more explosive movements such as a squat jump.

The small mean ranges across estimations based on different sets of input data and the good agreement between in vivo and estimated parameters further confirm the validity of the proposed approach to estimate calf muscle-tendon parameters.

In vivo and estimated gastrocnemius medialis optimal fiber lengths were in good agreement (in vivo young 47.5 ± 6.7 mm, in vivo older 44.0 ± 6.3 mm, estimated young 63.2 ± 20.7 mm, estimated older 46.9 ± 7.1 mm) ^18^. Also, in vivo and estimated gastrocnemius medialis optimal pennation angles were in good agreement (in vivo young 18.9 ± 2.4 °, in vivo older 18.4 ± 2.6 °, estimated young 17.6 ± 3.9 °, estimated older 16.4 ± 2.2) ^18^. We assume that we did not find any differences between young and older adults with respect to these parameters ^33^ due to the low sample size in this study. We observed good agreement between estimated gastrocnemius lateralis optimal fiber length and in vivo gastrocnemius lateralis fiber lengths during walking reported in other studies (in vivo young 68 ± 14 mm, in vivo older 63 ± 8 mm, estimated young 66.7 ± 22.0 mm, estimated older 49.6 ± 7.5 mm) ^23^. We could not compare in vivo and estimated tendon slack length because the in vivo free tendon length and in vivo aponeurosis length are both represented in the estimated tendon slack length in the Hill-type muscle model that is used in our musculoskeletal models ^36^. We found a larger mean range across estimations for Achilles tendon stiffness compared to other parameters. This probably relates to the differences between the in vivo Achilles tendon force-elongation behavior that is characterized by a toe region and the linear Achilles tendon force-elongation behavior without a toe region in our models. For instance, lower estimated (linear) Achilles tendon stiffness will better fit in vivo (with a toe region, curvilinear) force-elongation data at lower Achilles tendon forces during slower walking. Similarly, higher estimated Achilles tendon stiffness will better predict in vivo force-elongation data at higher Achilles tendon forces during faster walking. Indeed, Achilles tendon stiffness estimated at the lower speed (speed 1) was on average 15% lower than Achilles tendon stiffness estimated at the higher speed (speed 3). This explains the larger mean range across estimations of Achilles tendon stiffness compared to other parameters. We found good agreement between in vivo and estimated Achilles tendon stiffness estimated at speed 1 (in vivo young 170 ± 37 N/mm, in vivo older 141 ± 48 N/mm, estimated young 193.4 ± 53.3 N/mm, estimated older 183.0 ± 50.0 N/mm) ^33^. Therefore, we think that the use of a non-linear instead of linear model of the tendon force-length relation might further improve the results of this study ^22^.

The use of experimental ultrasound images and a simple Hill-type muscle models limits the accuracy of the parameter estimation. First, our results are dependent on the quality of the collected ultrasound data. The use of two-dimensional images to derive the length of muscle fibers that are not constrained to move in one plane results in measurement errors. To limit such errors, extreme care was taken to align the probe with the muscle fibers and the aponeuroses as described in previous literature ^3^. Moreover, echo intensity is higher in older adults due to adipose infiltration and connective tissue changes ^32^. This negatively influences image and image processing quality, which might explain the outlier in the analyses of the R^2^ between measured and generic or individualized gastrocnemius medialis fiber length (Fig. 2A). To limit the sensitivity of our approach to measurement errors, parameter estimation was based on data from multiple strides. Additionally, a systematic analysis defined the weights attributed to the different terms in the cost function to make sure that tracking the ultrasound measurements did not result in unrealistic muscle activations. Second, the Hill-type muscle model that was used in this study ^36^ represents the muscle-tendon unit by a series of line segments that do not capture the three-dimensional shape of the muscle. These three-dimensional shapes undergo changes during muscle contraction that can alter muscle-tendon interaction ^1^. The Hill-type muscle model also combines the characteristics of the free tendon and the elastic elements inside the muscle belly into one uniform tendon for every muscle. As such, it simplifies the non-uniform Achilles tendon dynamics ^9,10^. Moreover, our ultrasound data suggests that the assumption of constant muscle thickness is a strong simplification of the complex dynamics in the gastrocnemius medialis during walking. These simplifications along with the use of a linear tendon might explain the decreased fit between measured and calculated gastrocnemius medialis fiber length and pennation angle at the end of the push-off phase and during swing phase observed in some subjects (Fig. 3H and 3I). Because ankle moments and corresponding muscle forces are very low during these phases (Fig. 3J), we do not expect this misfit to influence the estimation of the metabolic cost a lot.

In conclusion, we proposed an approach to estimate calf muscle-tendon properties based on ultrasound data and muscle EMG collected during walking that improved calculations of calf muscle fiber lengths and activations. Moreover, validations at walking speeds that were not used for estimation and good agreement between estimated parameters and in vivo measured parameters from literature underline the accuracy of the new method. Calculations of calf muscle metabolic energy consumption highlight the importance of individualizing calf muscle-tendon parameters when investigating calf muscle-tendon interactions during walking.

